# Drinking motives and alcohol use among college students with the alcohol flush reaction

**DOI:** 10.1101/436469

**Authors:** Sarah Soyeon Oh, Yeong Jun Ju, San Lee, Sung-in Jang, Eun-Cheol Park

**Author notes:** Corresponding author: Eun-Cheol Park, MD, PhD, Department of Preventive Medicine and Institute of Health Services Research, Yonsei University College of Medicine, 50 Yonsei-ro, Seodaemun-gu, Seoul 120-752, Korea.

## Abstract

**Background:** This study investigated the relationship between drinking motives and alcohol use among a nationally representative sample of college students with the alcohol flush reaction (AFR). We surveyed and analyzed the data of 2,245 male and 2,326 female college students in a nationally representative sample of 82 colleges in South Korea.

**Methods:** Of our study population, 725 males (32.3%) and 812 females (34.9%) reported to currently suffering from AFR. Multiple regression analysis was used to identify the association between drinking motives and drinking behavior, measured via the AUDIT.

**Results:** Relative to drinking because of peer pressure, students drinking for pleasure (males: β = 2.622, *p* <.0001; females β = 2.769, *p* <.0001) or stress/depression (males: β = 2.479, *p* <.0001; females β = 2.489, *p* <.0001) had higher AUDIT scores. Among students drinking because of stress/depression, seniors (males: β = 3.603, *p*<.0001; females: β = 3.791, *p* = 0.000), smokers (males: β = 1.564, *p* = 0.000; females β = 1.816, *p* = 0.007) and/or liberal arts students (males: β = 6.1136, *p*<.0001; females β = 4.2105, *p* <.0001) consumed more alcohol than their peers. Relative to conformity motives, enhancement and coping motives were found to have a greater influence on alcohol intake among college students with alcohol flush reaction.

**Conclusion:** Considering that the flush reaction can occur in AFR individuals after just one sip of wine, our results show that educators and policymakers must take action to deal with this problem.

## INTRODUCTION

The alcohol flush reaction (AFR) is a symptom that approximately 36% of East Asians (Japanese, Chinese, and Koreans) have (Brooks et al., 2009). Also known as the “Asian Glow” or “alcohol-induced facial flushing”, individuals with this symptom are likely to experience a reddening of the face, nausea, and increased heart rate upon consuming alcohol (Ma et al., 2017).

AFR occurs when an individual inherits a deficiency in the single nucleotide polymorphism, ALDH2*504lys (ALDH2), which results in accretion of toxic acetaldehydes in the body (Adams and Rans, 2013). For individuals with AFR, toxic acetaldehyde build-up can occur from the first sip of alcohol (Price et al., 2004) which is extremely dangerous as acetaldehyde build-up has been classified as a Group 1 human carcinogen by the International Agency for Research on Cancer of the World Health Organization (Salaspuro, 2011). Low-activity ALDH2 has been uniformly associated with various cancers including esophageal (Brooks et al., 2009, Cui et al., 2009, Gu et al., 2012, Liu et al., 2017, Matsuo et al., 2001, Wu et al., 2013, Yang et al., 2007, Yang et al., 2010, Yu et al., 2018, Zhang et al., 2010), gastric (Chen et al., 2016, Duell et al., 2012, Ghosh et al., 2017, Hidaka et al., 2015, Ishioka et al., 2018, Jiang et al., 2017, Wang et al., 2014, Yang et al., 2017), colorectal (Crous-Bou et al., 2013, Guo et al., 2013, Matsuo et al., 2006, Xinhua and Yanfei, 2017, Yang et al., 2009, Zhao et al., 2016, Zhong et al., 2016), and stomach (Matsuo et al., 2013).

The association between AFR and cancer in the oral cavity and esophagus (squamous cell carcinoma) is particularly alarming as esophageal cancer has one of the lowest 5-year survival rates of all cancers in Korea for both males (35.0%) and females (37.3%) (Jung et al., 2017). Researchers have stated that if moderate or heavy drinking ALDH2 heterozygotes decreased their alcohol intake, more than 50% of esophageal squamous cell carcinomas could be prevented in East Asian populations (Brooks et al., 2009).

Unfortunately, alcohol-induced facial flushing is a research area that has been limitedly researched for a nationally representative sample of South Koreans, despite affecting around 1/3 of the population. This is alarming as 50,100 males and 30,500 females in Korea die from cancer annually, and the alcohol consumption (in liters of pure alcohol) per capita (15+) in Korea averages 12.3 liters, which is more than double the Western Pacific Region’s average of 5.4 liters. It is also unusual as ALDH2 deficiency is relatively simple for clinicians to determine; because of the intensity of symptoms, most flushers are able to answer to questions that determine ALDH2 deficiency (Brooks et al., 2009).

Furthermore, limited research has been done on what drinking motives influence individuals with alcohol-induced facial flushing to drink. Currently, there is only a consensus among academics that individuals with the reaction are less likely to drink than non-flushers with respect to both frequency and amount (Suwaki and Ohara, 1985, Suzuki et al., 1997). Research indicates that social motives, characterized by positive reinforcement generated externally from drinking alcohol with others, and enhancement motives, generated from the psychoactive effects of consuming alcohol, are the most common motives for alcohol use among university students (Maphisa, 2018 #8). Thus, given the limited literature on drinking motives among individuals with AFR, two important questions remain: 1. Do the same drinking motives apply to college students, who suffer from the irritating symptoms of AFR? 2. Which drinking motives increase alcohol consumption, and the risk of alcohol-related problems among individuals with AFR? Therefore, the present study focuses on examining the association between drinking motives and alcohol use among individuals with AFR in South Korea. It also investigates the factors that strengthen or discourage this relationship.

## MATERIALS AND METHODS

### Study population and data

In the 2017 national statistics published by the Korean Educational Development Institute on college students, we found that 1,951,940 students (4-year: 1,506,745; liberal arts: 445,195) are enrolled in 356 colleges (4-year: 195, liberal arts: 161) in South Korea. Thus, we stratified a proportionately representative sample of undergraduate students from 54 4-year colleges and 28 liberal arts colleges. Students in these colleges were further stratified according to major, sex, and region (Seoul, Incheon/Gyeonggi, Gangwon, Daejeon/Chungjeong, Gwangju/Jeolla, Daegu/Gyeongbuk, Busan/Ulsan/Gyeongnam) to be representative of national statistics. After completion of a small pilot study, we randomly selected 4,571 students from this sample for our questionnaire. Of these students, we extracted the data of those who self-reported to currently suffering from alcohol-induced facial flushing, for a final study population of 725 males and 812 females with AFR.

Data was collected via face-to-face surveys with students. The 14-page survey instrument asked students a number of questions about their drinking behavior, health, and thoughts on campus-alcohol policy. Whenever possible, the instrument included alcohol-related questions that had been previously used in other international, national or large-scale epidemiological studies including the Harvard College Alcohol Study (Wechsler et al., 2002), the Korea National Health and Nutrition Examination Survey (KNHANES), and the Korea Youth Risk Behavior Web-Based Survey (KYRBS).

A standard drink was defined as the amount of alcohol contained in one glass of alcohol drink (approximately 8 grams of pure alcohol), equivalent to:1 shot of soju, 1 glass of bottled beer, 2/3 of a canned beer, 1/2 glass of draft beer, 1/2 bowl of makgeolli (rice wine), 1/2 glass of wine, 1 glass of whiskey, 1 shot of cheongju (refined rice wine), 1 shot of herbal liquor, 1 shot of fruit wine, or a 3/5 glass of mixed liquor (soju+beer), in accordance with the standards of the Korea Centers for Disease Control & Prevention.

## Measures

### Outcome Variable

In this study, alcohol intake, measured through the Alcohol Use Disorders Identification Test (AUDIT), was selected as the outcome variable. The AUDIT questionnaire was given in its original format, consisting of ten questions related to frequency of drinking, number of drinks per session, frequency of heavy drinking, impaired control following drinking, morning-after drinking, and feelings of guilt upon drinking, frequency of blackouts, alcohol-related injuries, and family concern over drinking.

### Drinking Motives

Drinking motives were measured via individual answers to the question, “In the last 12 months, what was your main reason for drinking alcohol?” Answers were categorized into “peer pressure,” “pleasure,” “stress/depression,” “boredom,” and “other.” ‘Other’ responses included the following: “force of habit,” “birthday,” “anniversary,” “to sleep.”

### Statistical analysis

Frequencies and mean AUDIT scores were calculated for each variable through *t* tests and analyses of variance. To examine the relationship between reason for alcohol consumption and AUDIT score, multiple logistic regression analysis was performed, after controlling for the following confounders: year level, major, GPA, pocket money, living status, smoking status, stress level, depressive thoughts, suicidal thoughts, suicidal attempt, underage drinking experience, and number of sororities/clubs.

Alcohol-induced facial flushing was measured through the questionnaire created by Yokoyama and Omori, which asks the following questions to determine past and present ALDH2 heterozygotes: 1) Do you have a tendency to develop facial flushing immediately after drinking a glass (about 180 mL) of beers? 2) Did you have the tendency in the first one or two years after you started drinking? This questionnaire was rated to have a high sensitivity rate of 90.1% and specificity rate of 88.0% (Yokoyama et al., 2010) in classifying respondents into never, former, and current sufferers of ALDH2 deficiency.

The simultaneous relationship between alcohol use and other factors like year level and amount of pocket money, according to reason for alcohol consumption, were determined by subgroup analyses. The calculated p-values in this study were considered significant if lower than 0.05. All analyses were performed using SAS software, version 9.4 (SAS Institute, Cary, North Carolina, USA).

## RESULTS

Of the entire study population, the prevalence of alcohol-induced facial flushing was 32.3% for male students and 34.9% for female students. Mean AUDIT scores of students with AFR were 8.253 ± 5.341 for males and 7.233 ± 5.474 for females, whilst non-AFR students had mean AUDIT scores of 10.518 ± 5.379 and 10.369 ± 6.144, respectively. Relative to never- and past-sufferers of facial flushing, individuals with current facial flushing were less likely to have high AUDIT scores as illustrated in Figure 1.

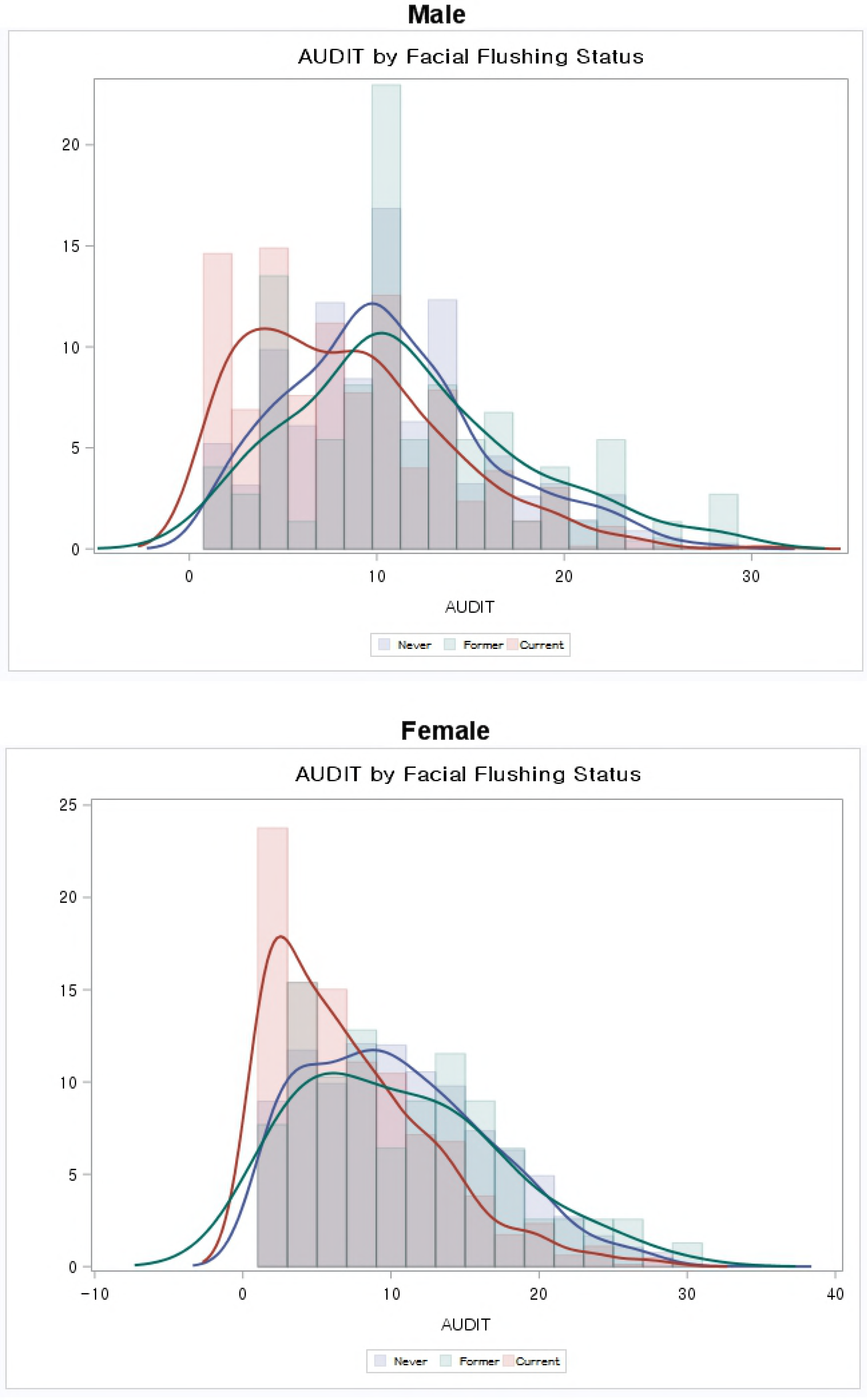

Table 1 shows the general characteristics of the AFR population. 34.5% of the male population (n 250) and 31.8% of the female population (n 258) reported to consuming alcohol because of peer pressure. This was followed by individuals who drink because of pleasure (males: 29.7%; females: 30.7%) or stress/depression (males: 22.5%; females: 22.5%). Individuals who consume alcohol because of pleasure had the highest AUDIT scores (males: 9.758 ± 4.938; females: 9.012 ± 5.487).

**Table 1.** General characteristics of ARF individuals and AUDIT scores.

|  | AUDIT Score |  |  |  |  |  |  |  |
| --- | --- | --- | --- | --- | --- | --- | --- | --- |
|  | Male Flushers (n 725) |  |  |  | Female Flushers (n 812) |  |  |  |
|  | N | Mean | SD | p-value | N | Mean | SD | p-value |
| **Drinking Motive** |  |  |  |  |  |  |  |  |
| Peer Pressure (Conformity) | 250 | 6.748 | 5.453 | <.0001 | 258 | 5.481 | 4.911 | <.0001 |
| Pleasure (Enhancement) | 215 | 9.758 | 4.938 |  | 249 | 9.012 | 5.487 |  |
| Stress/Depression (Coping) | 163 | 9.460 | 5.319 |  | 183 | 8.251 | 5.899 |  |
| Boredom | 87 | 6.874 | 4.600 |  | 111 | 6.036 | 4.375 |  |
| Other | 10 | 5.900 | 3.665 |  | 11 | 3.273 | 3.101 |  |
| **Year Level** |  |  |  |  |  |  |  |  |
| 1 | 224 | 8.049 | 5.454 | 0.397 | 259 | 7.201 | 5.464 | 0.002 |
| 2 | 215 | 8.656 | 5.383 |  | 255 | 7.859 | 5.629 |  |
| 3 | 128 | 8.500 | 5.073 |  | 149 | 7.705 | 5.587 |  |
| ≥4 | 158 | 7.797 | 5.336 |  | 149 | 5.752 | 4.842 |  |
| **Major** |  |  |  |  |  |  |  |  |
| Humanities and Social Sciences | 323 | 8.560 | 5.610 | 0.102 | 411 | 7.007 | 5.312 | 0.067 |
| Engineering / Natural Sciences | 343 | 7.828 | 5.004 |  | 281 | 7.822 | 5.799 |  |
| Liberal Arts | 59 | 9.051 | 5.609 |  | 120 | 6.633 | 5.151 |  |
| **GPA** |  |  |  |  |  |  |  |  |
| ≥4.0 | 117 | 7.940 | 5.081 | 0.113 | 122 | 6.705 | 4.825 | 0.000 |
| 3.5-4.0 | 238 | 8.176 | 5.376 |  | 321 | 6.408 | 5.056 |  |
| 3.0-3.5 | 261 | 7.992 | 5.239 |  | 275 | 8.044 | 5.934 |  |
| ≤3.0 | 109 | 9.385 | 5.696 |  | 94 | 8.372 | 5.773 |  |
| **Pocket Money** |  |  |  |  |  |  |  |  |
| Q1 (Low) | 276 | 7.315 | 4.885 | 0.000 | 320 | 5.844 | 4.602 | <.0001 |
| Q2 | 177 | 8.181 | 5.462 |  | 219 | 7.306 | 5.619 |  |
| Q3 | 140 | 9.243 | 5.818 |  | 141 | 8.064 | 5.388 |  |
| Q4 (High) | 132 | 9.265 | 5.258 |  | 132 | 9.598 | 6.266 |  |
| **Living Status** |  |  |  |  |  |  |  |  |
| Family Home | 347 | 7.755 | 5.178 | 0.006 | 467 | 6.478 | 5.193 | <.0001 |
| Living Alone/Flatting | 201 | 9.244 | 5.883 |  | 168 | 9.250 | 6.103 |  |
| College Dorm | 177 | 8.107 | 4.866 |  | 177 | 7.316 | 5.091 |  |
| **Smoking Status** |  |  |  |  |  |  |  |  |
| Current Smoker | 262 | 9.435 | 5.278 | <.0001 | 61 | 9.770 | 5.763 | <.0001 |
| Past Smoker | 64 | 10.047 | 5.411 |  | 22 | 12.682 | 7.247 |  |
| Non-Smoker | 399 | 7.190 | 5.141 |  | 729 | 6.857 | 5.249 |  |
| **Stress Level** |  |  |  |  |  |  |  |  |
| High | 74 | 8.824 | 4.793 | 0.448 | 121 | 7.471 | 6.122 | 0.410 |
| Normal | 506 | 8.281 | 5.504 |  | 574 | 7.308 | 5.406 |  |
| Low | 145 | 7.869 | 5.023 |  | 117 | 6.624 | 5.090 |  |
| **Depressive Thoughts** |  |  |  |  |  |  |  |  |
| Yes | 73 | 9.753 | 5.459 | 0.011 | 133 | 8.662 | 6.423 | 0.001 |
| No | 652 | 8.086 | 5.306 |  | 679 | 6.954 | 5.228 |  |
| **Suicidal Thoughts** |  |  |  |  |  |  |  |  |
| Yes | 18 | 11.778 | 6.320 | 0.005 | 35 | 9.486 | 7.751 | 0.013 |
| No | 707 | 8.164 | 5.289 |  | 777 | 7.133 | 5.333 |  |
| **Suicidal Attempt** |  |  |  |  |  |  |  |  |
| Yes | 9 | 16.111 | 5.487 | <.0001 | 14 | 11.500 | 8.636 | 0.003 |
| No | 716 | 8.155 | 5.269 |  | 798 | 7.159 | 5.380 |  |
| **Underage Drinking Experience** |  |  |  |  |  |  |  |  |
| Yes | 397 | 9.008 | 5.380 | <.0001 | 355 | 8.225 | 5.811 | <.0001 |
| No | 328 | 7.341 | 5.156 |  | 457 | 6.464 | 5.071 |  |
| **Number of sororities/clubs** |  |  |  |  |  |  |  |  |
| None | 345 | 7.710 | 5.124 | 0.005 | 422 | 6.882 | 5.305 | 0.008 |
| One | 303 | 8.479 | 5.271 |  | 308 | 7.263 | 5.354 |  |
| Two or more | 77 | 9.805 | 6.211 |  | 82 | 8.939 | 6.445 |  |
| **Total** | **725** | **8.253** | **5.341** |  | **812** | **7.233** | **5.474** |  |

Table 2 shows the results of multiple regression analysis performed to investigate the relationship between various factors and AUDIT score among college students with alcohol-induced facial flushing. Certain drinking motives had a statistically significant association with increased alcohol intake: relative to drinking because of peer pressure, students drinking for pleasure (males: β= 2.622, *p* <.0001; females β = 2.769, *p* <.0001) or stress/depression (males: β = 2.479, *p* <.0001; females β = 2.489, *p* <.0001) had higher AUDIT scores. Whilst year level had no statistically significant association with alcohol intake for male flushers, female flushers in their senior year or above scored lower on the AUDIT than those in their freshman year (β = −1.587, *p* = 0.002). Students receiving higher amounts of pocket money were likely to drink more than students with lower levels of income (males: β = 1.093, *p* = 0.039; females β = 12.739, *p* <.0001). For both males and females, relative to non-smokers, past (males: β = 1.994, *p* = 0.003; females β = 5.090, *p* <.0001) and current smokers (males: β = 1.564, *p* = 0.000; females β = 1.816, *p* = 0.007) were likely to score higher on the AUDIT, as well as females with underage drinking experience (β = 0.714, *p* = 0.042) compared to those without such experience. For both males and females, those in two or more sororities/clubs had higher levels of alcohol intake than those in no sororities/clubs (males: β = 1.383, *p* = 0.030; females β = 1.156, *p* = 0.052).

**Table 2.** Results of the GEE analyzing Reason for Alcohol Consumption and AUDIT of ARF individuals.

|  | AUDIT Score |  |  |  |  |  |
| --- | --- | --- | --- | --- | --- | --- |
|  | Male Flushers |  |  | Female Flushers |  |  |
| | $\beta^*$ | S.E | p-value | $\beta^*$ | S.E | p-value |
| **Drinking Motive** |  |  |  |  |  |  |
| Peer Pressure | Ref. |  |  |  |  |  |
| Pleasure | 2.622 | 0.460 | <.0001 | 2.769 | 0.437 | <.0001 |
| Stress/Depression | 2.479 | 0.503 | <.0001 | 2.489 | 0.475 | <.0001 |
| Boredom | -0.261 | 0.607 | 0.667 | 0.469 | 0.551 | 0.394 |
| Other | -0.785 | 1.565 | 0.616 | -2.031 | 1.479 | 0.170 |
| **Year Level** |  |  |  |  |  |  |
| 1 | Ref. |  |  | Ref. |  |  |
| 2 | 0.735 | 0.468 | 0.116 | 0.344 | 0.427 | 0.420 |
| 3 | 0.042 | 0.556 | 0.940 | 0.065 | 0.503 | 0.898 |
| $\geq 4$ | -0.425 | 0.526 | 0.419 | -1.587 | 0.508 | 0.002 |
| **Major** |  |  |  |  |  |  |
| Humanities and Social Sciences | Ref. |  |  | Ref. |  |  |
| Engineering / Natural Sciences | -0.421 | 0.383 | 0.272 | 0.512 | 0.375 | 0.173 |
| Liberal Arts | 0.341 | 0.692 | 0.622 | -0.472 | 0.509 | 0.354 |
| **GPA** |  |  |  |  |  |  |
| $\geq 4.0$ | Ref. | | | Ref. | | |
| 3.5-4.0 | -0.121 | 0.555 | 0.827 | 0.014 | 0.519 | 0.978 |
| 3.0-3.5 | -0.003 | 0.541 | 0.996 | 1.334 | 0.528 | 0.012 |
| $\leq 3.0$ | 1.334 | 0.654 | 0.042 | 0.805 | 0.666 | 0.227 |
| **Pocket Money** |  |  |  |  |  |  |
| Q1 (Low) | Ref. |  |  | Ref. |  |  |
| Q2 | 0.485 | 0.473 | 0.305 | 1.161 | 0.429 | 0.007 |
| Q3 | 1.358 | 0.522 | 0.009 | 1.874 | 0.493 | 0.000 |
| Q4 (High) | 1.093 | 0.530 | 0.039 | 2.739 | 0.514 | <.0001 |
| **Living Status** |  |  |  |  |  |  |
| Family Home | Ref. |  |  | Ref. |  |  |
| Living Alone/Flatting | 0.842 | 0.445 | 0.059 | 1.635 | 0.446 | 0.000 |
| College Dorm |  |  |  |  |  |  |
| **Smoking Status** |  |  |  |  |  |  |
| Current Smoker | 1.564 | 0.407 | 0.000 | 1.816 | 0.667 | 0.007 |
| Past Smoker | 1.994 | 0.670 | 0.003 | 5.090 | 1.056 | <.0001 |
| Non-Smoker | Ref. |  |  | Ref. |  |  |
| **Stress Level** |  |  |  |  |  |  |
| High | 0.463 | 0.736 | 0.529 | -0.876 | 0.685 | 0.201 |
| Normal | 0.227 | 0.464 | 0.625 | 0.110 | 0.494 | 0.824 |
| Low | Ref. |  |  | Ref. |  |  |
| **Depressive Thoughts** |  |  |  |  |  |  |
| Yes | 0.521 | 0.646 | 0.419 | 0.347 | 0.524 | 0.508 |
| No | Ref. |  |  | Ref. |  |  |
| **Suicidal Thoughts** |  |  |  |  |  |  |
| Yes | 1.632 | 1.236 | 0.186 | 0.164 | 1.026 | 0.873 |
| No | Ref. |  |  | Ref. |  |  |
| **Suicidal Attempt** |  |  |  |  |  |  |
| Yes | 6.057 | 1.700 | 0.000 | 3.241 | 1.495 | 0.030 |
| No | Ref. |  |  | Ref. |  |  |
| **Underage Drinking Experience** |  |  |  |  |  |  |
| Yes | 0.645 | 0.378 | 0.088 | 0.714 | 0.351 | 0.042 |
| No | Ref. |  |  | Ref. |  |  |
| **Number of sororities/clubs** |  |  |  |  |  |  |
| None | Ref. |  |  | Ref. |  |  |
| One | 0.460 | 0.392 | 0.240 | 0.255 | 0.369 | 0.489 |
| Two or more | 1.383 | 0.637 | 0.030 | 1.156 | 0.594 | 0.052 |

Figure 2 shows the results of the subgroup analysis for the association between reason for alcohol consumption and AUDIT scores by year level, major, and amount of pocket money. Regarding year level, for male flushers, alcohol intake was highest among freshmen consuming alcohol because of stress/depression (β = 4.213, *p*<.0001) and seniors consuming alcohol for pleasure (β = 3.603, *p*<.0001). For female flushers, alcohol intake was highest among sophomores (β = 4.101, *p*<.0001) and seniors (β = 3.791, *p* = 0.000) consuming alcohol for pleasure. Regarding major, for both sexes, alcohol intake was highest among liberal arts students consuming alcohol because of stress/depression (males: β = 6.1136, *p*<.0001; females β = 4.2105, *p* <.0001). Lastly, female students receiving high amounts of pocket money were likely to consume large amounts of alcohol for both pleasure (β = 5.454, *p*<.0001) and stress/depression-related (β = 5.833, *p* = 0.000) purposes, relative to students with low levels of pocket money.

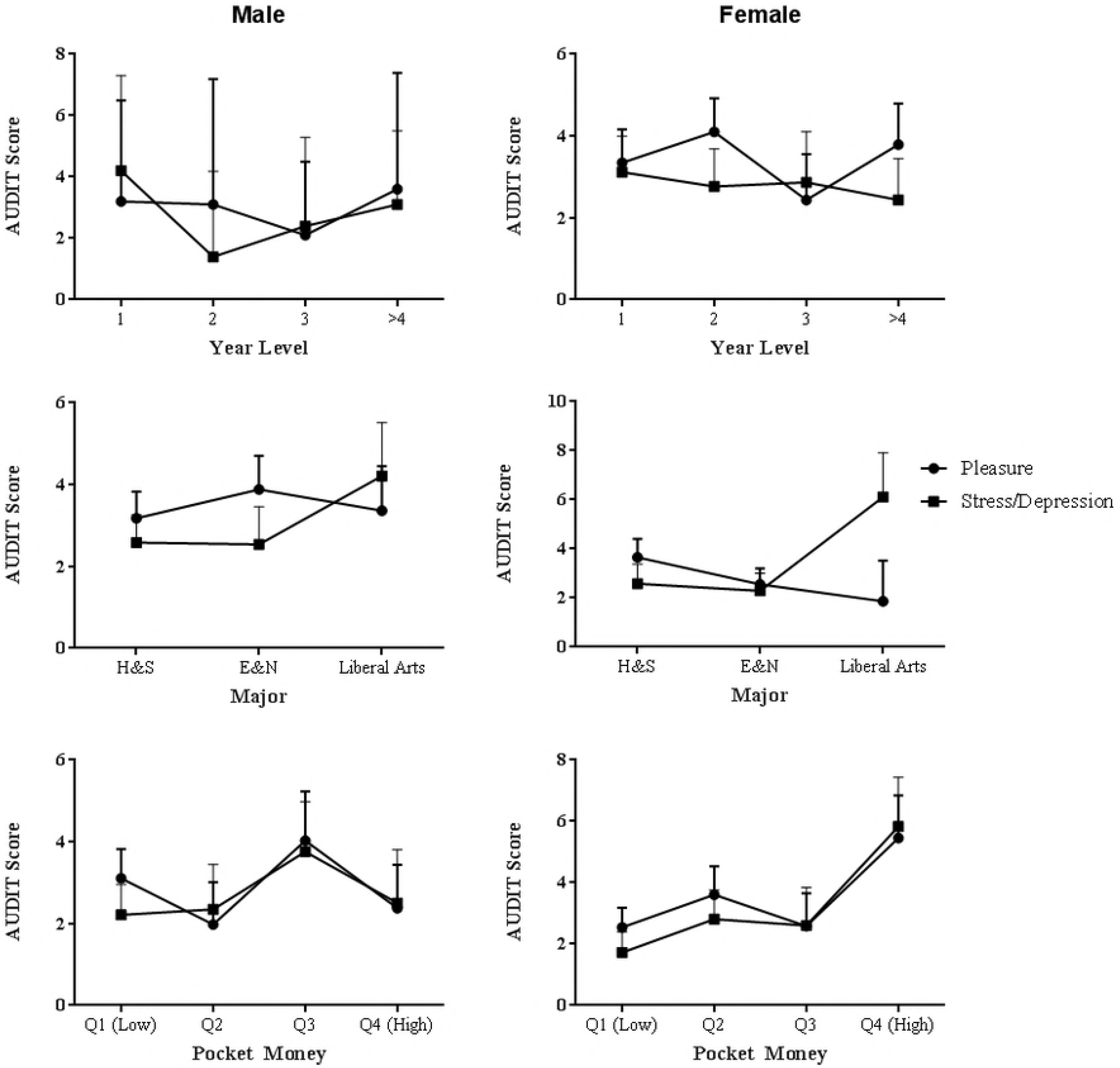

## DISCUSSION

In individuals with AFR, the flush-reaction can begin to occur after just one sip of wine (Price et al., 2004). Such individuals, who have an inability to metabolize the substance, should not be encouraged to consume any alcohol. However, our study shows that individuals with AFR, especially those who drink for pleasure or to alleviate stress/depression drink above this level. As seen in Table 1, the mean AUDIT score for flushers in our study population was 8.253 ± 5.341 for males and 7.233 ± 5.474 for females. This is above the AUDIT cut-off for hazardous drinking among normal populations without AFR of 10 (males) and 5 (females) (male cut-off: sensitivity, 81.90%; specificity, 81.33%; positive predictive value, 77.2%; negative predictive value, 85.3%; female cut-off sensitivity, 100.00%; specificity, 88.54%; positive predictive value, 52.6%; negative predictive value, 100.0%) (Chang et al., 2016). Female students with AFR especially, drink well-above this cut-off level, especially if their drinking motive is “pleasure,” “stress/depression,” or “boredom.”

Furthermore, our study reveals what the existing body of literature has failed or refused to acknowledge: individuals with AFR often enjoy alcohol consumption and this is both hazardous and dangerous for their health. Because of the irritating symptoms of AFR, acetaldehyde dehydrogenase inhibition has been used by drug-makers to develop effective treatments for alcoholics (Brewer et al., 2017, Ulrichsen et al., 2010). However, these symptoms have clearly not been powerful enough to prevent the hazardous drinking of around 30% of AFR individuals in our study population, who find drinking pleasurable.

Lastly, our research shows that the drinking motives behind college students with AFR slightly differ from the drinking motives of college students without AFR. The existing body of research states that social and enhancement motives are the most frequently endorsed by university students, whilst coping motives are found to be most strongly associated with negative alcohol consequences (Merrill, 2010 #10). However, our study reveals that among college students with AFR, enhancement motives like pleasure, are the strongest drinking motive and most likely to cause risk of Alcohol Use Disorder, followed closely by coping motives like stress/depression.

This study has several limitations that should be considered when interpreting results. First, our study is cross-sectional in design; thus, caution should be exercised in interpreting causality between reason for alcohol consumption and alcohol intake. Furthermore, the Yokoyama and Omori questionnaire for determining ALDH2 deficiency, as well as the AUDIT cut-off values set by Chang, are limited in reliability and validity relative to clinical evaluations of these symptoms by a healthcare professional.

Second, there are not enough previous studies with regard to a nationally representative population of Koreans when it comes to measuring alcohol-induced facial flushing and its effect on drinking behavior/related problems. It is difficult to see whether the values we calculated are similar to that of the statistics found in previous studies for Koreans, especially for the college students” age group. Third, various biases may have emerged from our sampling and surveying methods; because college students in South Korea drink large amounts of alcohol relative to adults, different patterns are likely to emerge in an adult sample. Likewise, a small number of Christian colleges that were originally in our sample declined our request for participation because of their teetotalism principles and thus, had to be replaced with non-Christian colleges. Because of the face-to-face method that we employed for accuracy of obtaining responses to complicated questions, there may have been response biases, relative to social desirability. The majority of questions in our survey instrument required students to think about their drinking behaviors in the last 12 months or so, which likely resulted in recall bias. Finally, although we included numerous lifestyle covariates as potential confounders, the limited nature and number of questions in our instrument made it difficult for other confounding variables, relative to health, socio-demographics, gene-environment, and lifestyle, to be measured and controlled.

Despite these limitations, our study also has several strengths. Few studies have measured ALDH2 deficiency for a nationally representative population in South Korea, and fewer studies have taken an epidemiological approach to see the association between ALDH2 deficiency and relative change in drinking behavior, especially with regard to the reasons that encourage individuals with AFR to drink, despite its irritating symptoms. Our study has found certain risk-groups in the college population that overly drink despite suffering from the flush reaction. Researchers, educators, and policy-makers are encouraged to further investigate, and target such students when creating and influencing campus alcohol policy.

